# Optimizing T-cell-mediated cancer killing: An agent-based model

**DOI:** 10.1101/2025.10.21.683801

**Authors:** Andrew P. Yu, Xingjian Zhang, Lance L. Munn

**Affiliations:** Edwin L. Steele Laboratories, Department of Radiation Oncology, Massachusetts General Hospital Cancer Center, Massachusetts General Hospital and Harvard Medical School, Boston, Massachusetts, United States of America; Boston Latin School, Boston, Massachusetts, United States of America

**Keywords:** Agent-Based Model, T-cell-Mediated Cytotoxicity, Optimization

## Abstract

Immunotherapies for cancer aim to improve immune cell function by helping immune cells recognize and kill tumors. A group of immunotherapies target T-cells, aiming to improve the patient’s prognosis. However, these therapies are not always successful, and the underlying mechanisms of T-cell elimination of tumors remain poorly understood. We developed a three-dimensional agent-based model to explore this behavior that probabilistically models cell growth, migration and interaction parameters. Through this project, we explore the influence of cytokines like IL-2 and CCL2 on T-cell behavior and optimize the interactions between T-cells and cancer cells to most efficiently kill tumors using modeling. How individual parameters such as T-cell killing rate, proliferation, infiltration and cytotoxicity is still largely unexplored, and our study aims to fill that gap by stochastically simulating T-cells as they attack a cancer spheroid in order to identify new potential strategies to enhance T-cell killing of tumors. Our study found that, from greatest to least effect, increasing the killing, proliferation, and infiltration rate of T-cells increased the clearance rate of the tumor. A 17% increase in killing rate was enough to control and shrink the tumor, while a 15% increase in their proliferation rate was sufficient to control the tumor without regressing it; a 30% increase in their proliferation rate was sufficient to shrink the tumor. Increasing the infiltration was only able to control the tumor, not cause it to regress.

## Introduction

Cancer remains one of the leading causes of death worldwide, with 9.7 million cancer-related deaths worldwide and nearly 20 million diagnoses in 2022 (1). Immunotherapies revolutionized the cancer treatment landscape, but even with countless new emerging therapies, the exact mechanism of cell-cell interactions between cancer cells and T-cells remains poorly understood (2). Computational models have begun to rise in popularity as they have proven useful for studying tumor behavior and therapeutic options (3–5). Investigations into checkpoint blockades and other therapeutic interventions have been conducted (6), and we now apply these approaches to study T-cell cytotoxicity as dictated by different therapies to improve tumor elimination and evaluate their efficacy. We investigated and tuned the model in the context of triple negative breast cancer. We use two known cytokines produced by cancer cells and T-cells, respectively: CCL2 and IL-2, respectively.

IL-2 is known to support T-cell proliferation and cancer killing (7), and is produced by activated CD8^+^ T-cells (8). Additionally, it acts as a T-cell chemoattractant (9), Specifically, it acts on the tumor microenvironment and improves activation of various cell types present (8,10). On the other hand, cancer is known to secrete CCL2 to drive proliferation of supporting immunosuppressive cells such as fibroblasts and Regulatory T-cells (11,12). CCL2 is also a potent T-cell chemo-attractant (9), leading to its incorporation into this simulation.

This simulation seeks to identify the effectiveness of therapeutic methods developed to enhance T-cell function by isolating each variable individually. Our hypothesis was first that, without any treatment interventions, T cells would be unable to control the tumor. To recapitulate current therapeutic strategies on altering T cell behaviors, we modulate T cell Killing Rate, Proliferation, and Movement Speed to investigate their respective effectiveness on increasing the efficiency of T-cell mediated tumor control. Then, by increasing each variable in isolation, we were able to identify new behavior that affects the T cells’ ability to kill.

We performed *in silico* experiments in MATLAB with an agent-based model of T-cells and cancer cells with stochastic behavior. The *in-silico* nature of the simulations allows for the isolation and precise control of each variable, and then for the testing of multiple variables at the same time in multi-modal treatments. These cells interact in three dimensional matrices and are bounded by rules including cell division, migration, and killing, with underlying cytokine matrices determining their behavior. The simulation layers are summarized in Figure 1a, which shows blue T cells and red cancer cells on the left and a 3-dimensional heat-map for the middle *x-y* slice of an example CCL2 matrix on the right. As seen in Figure 3b, cancer cells and T-cells are able to move within a radius around themselves in their respective matrices in three dimensions to any space unoccupied by other cells. Because each cancer cell and T cell acts independently every timestep, the model is agent-based. The full flow of the model as each timestep is processed is discussed in the Methods section.

**Figure 1:**
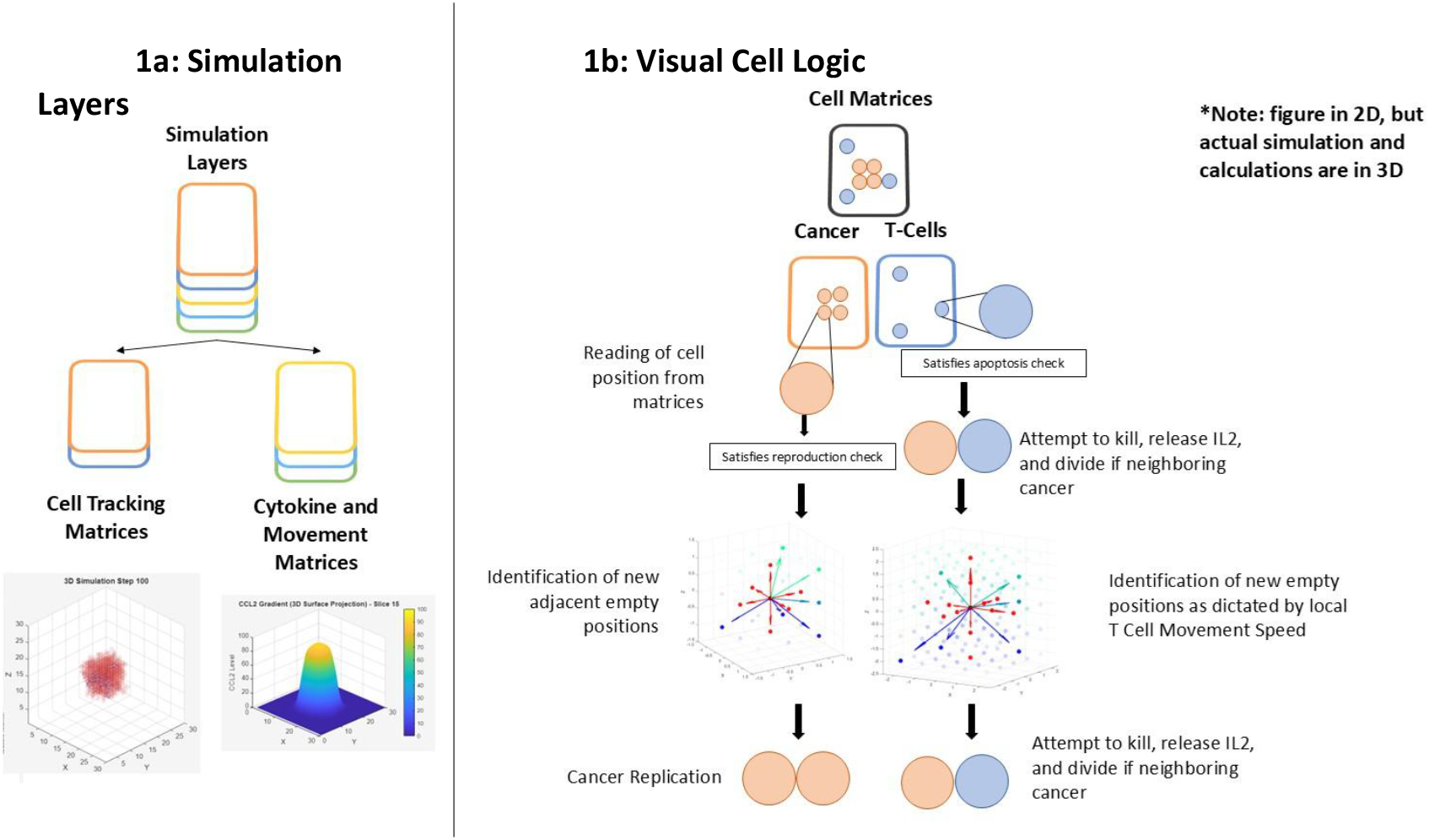
Simulation Overview.

**Figure 2:**
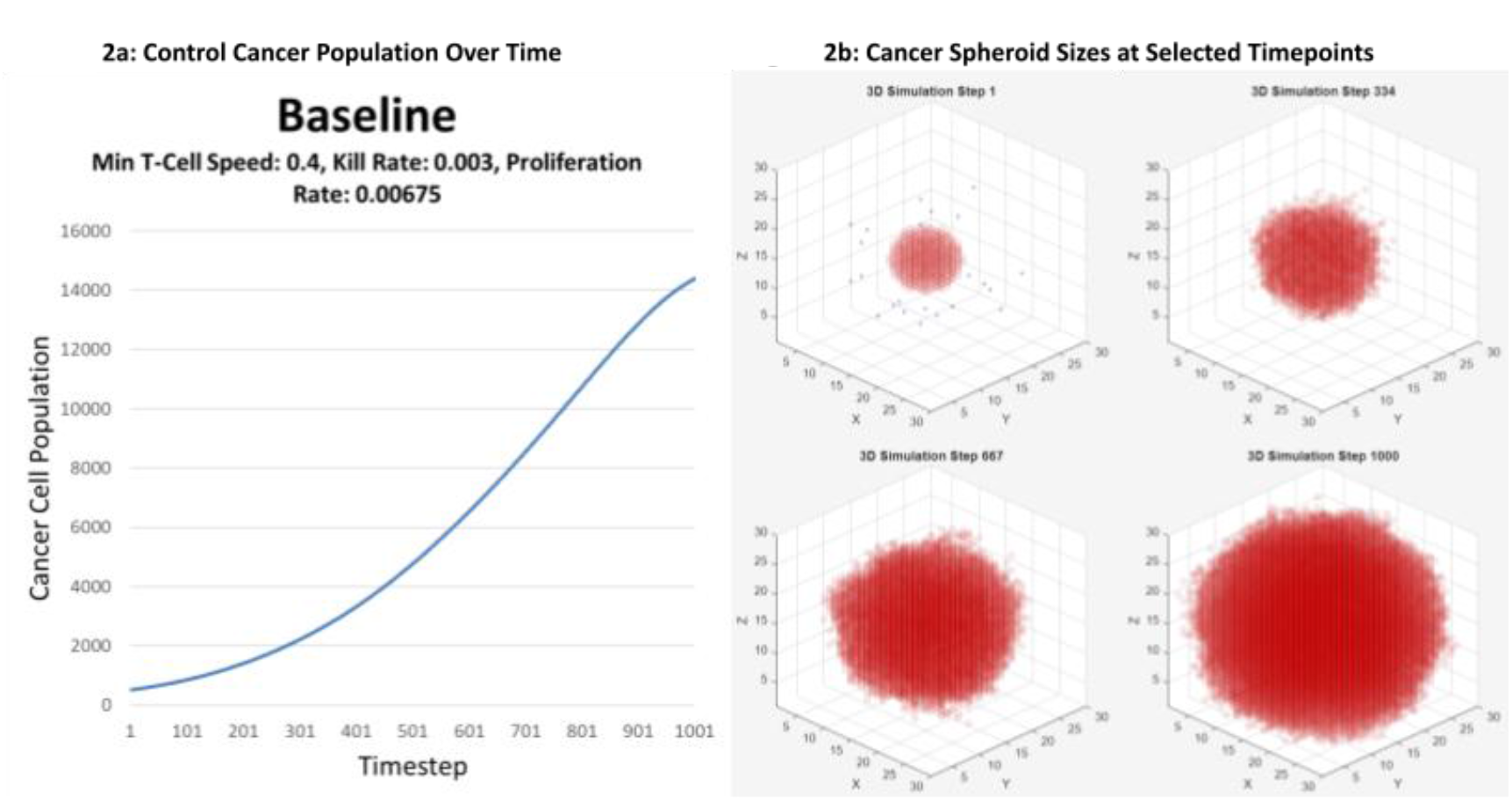
Control Condition Plots

**Figure 3:**
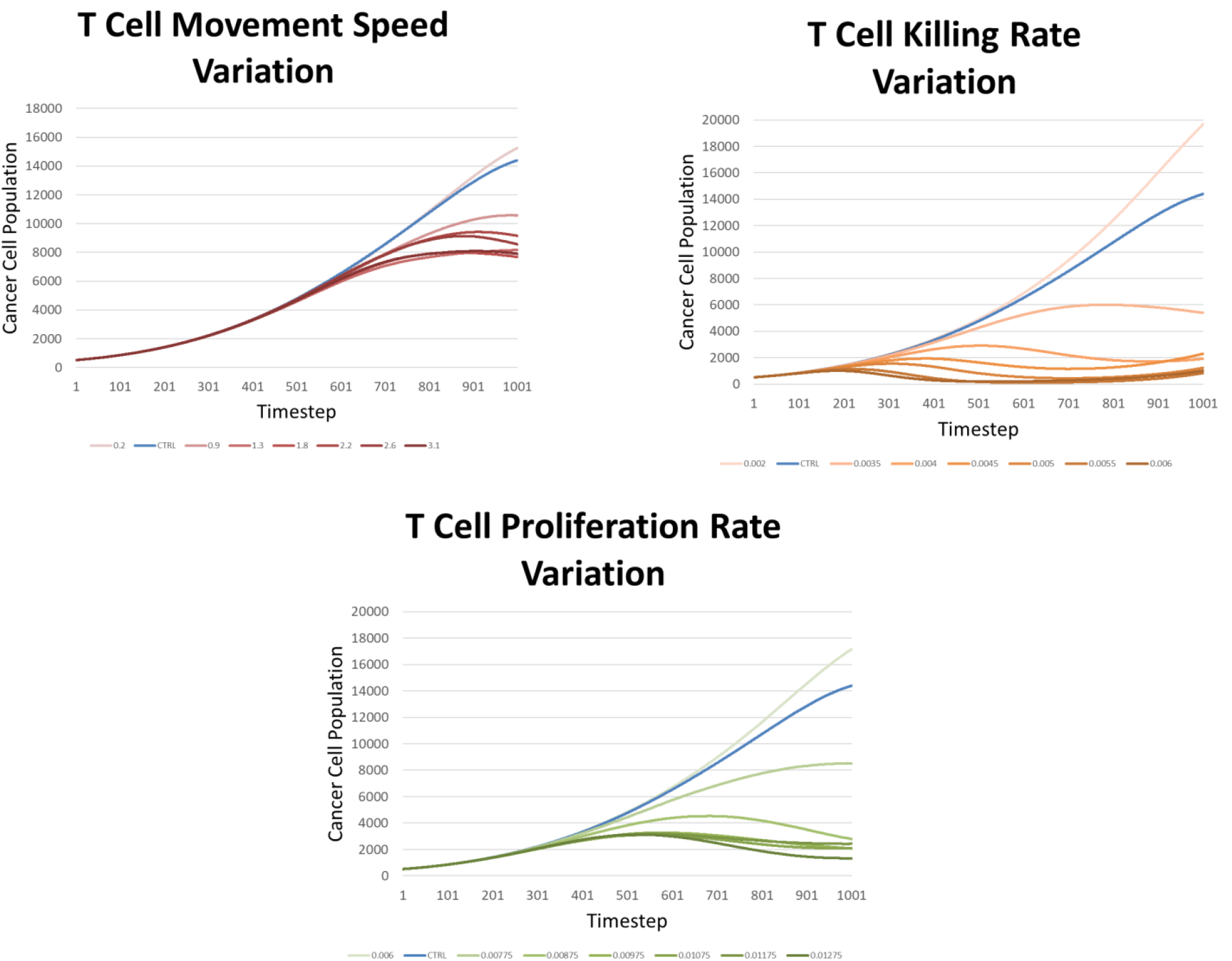
Average Cancer Population over Time for Varied Parameter

## Results

As a control, 25 T-cells were seeded with the tumor with known untreated tumor conditions: 520 tumor cells, a Min-T-cell Speed of 0.4 Units/Timestep (which matches clinical observation of T cell movement being 1.36 μm/min (13)), a killing rate of 0.003, and a proliferation chance of 0.00675 (these values were obtained by manually analyzing time-lapse footage of T-cells against cancer). With these conditions, T-cells were not able to control the tumor, with the tumor growth slowing only as the tumor exceeded the bounds of the simulation space. 100 simulations under those conditions were performed, with the mean displayed below:

Subsequently, 100 trials were conducted for each combination of presets, varying each parameter individually up to double its value in increments of 5% and including one value below to test the sensitivity.

The means for each group of simulations are summarized below, with the blue line being the same control group, and darker lines indicating an increase in each parameter:

For the tumor movement speed group, each increasing speed value decreased the end tumor population, approaching a reduction of 50%. Interestingly, the T cells are not able to regress the tumor with this treatment and are only able to moderately control its growth (the population does not decrease significantly for most simulations).

On the other hand, increasing the T-cell Killing rates was the most effective, with just a 10% increase in the killing rate causing control of the tumor population and further increase leading to regression of the tumor. It should be noted, however, that much of the time, the T cells were not able to completely eradicate the cancer and after about 100 timesteps with a low population, the cancer population would rebound. In these simulations, much of the time, the T cells would eradicate the main tumor spheroid in the center of the simulation, but a few cancer cells would escape to the corners of the simulation, where, because of the T cells’ slow speed, the T cells are unable to follow. This is compounded by the relatively lower density of cancer (causing a lower CCL2 concentration) and the lack of T cell killing (causing a lower IL2 concentration) on the edges of the simulation, which decreases T-cell movement towards areas until the cancer population grows out of control.

When the T-cell proliferation was varied, the cancer populations decreased to a degree between the killing rate and tumor movement speed groups. For most of the curves, moreover, even for controlled tumors, it is apparent that the cancer population is rebounding.

In order to quantitatively understand the implications of this, the area under the curve of the cancer population across time (henceforth referred to as the AUC) was calculated for each simulation, and box plots were created to summarize each parameter. The box plots are plotted on top of violin plots, which show the shape of the distributions. Then, a 1-way ANOVA test was conducted for each condition against the control group, with significant results for each group. Finally, a Dunnett’s test was performed comparing each group to the control and stars added according to the result (1 star for *p*<0.05, 2 stars for *p*<0.01, and 3 stars for *p*<0.01).

The T-cell Movement Speed Variation shows a decrease in the mean and median for all increases in speed, but the distribution is wide, showing that the simulations were highly heterogeneous and the T cells were not always able to control the cancer. The local minima of Minimum Speeds of 1.8 and 2.3 validate for the first time the theory that increased T-cell movement may not always lead to a direct increase in tumor reduction. In this case, the intermediate T-cell Speeds were able to penetrate the tumor but were unable to move outwards fast enough to curb the tumor shell’s expansion.

The Killing Rate plots show a marked decrease in both the spread of the AUCs and the values of the AUCs themselves, indicating that this therapeutic mode is highly effective at controlling the tumor population. The proliferation rate plots demonstrate that its effectiveness lies between the speed variation and the killing rate variation.

Interestingly, the violin plots show that there are several simulations for every group that have areas under the curve of about 7*10^6^ units, which correspond to simulations where the T cells were completely unable to penetrate into the tumor at all. Without IL2 released by tumor killing, the T cells quickly apoptosed, so the cancer was able to divide uncontrollably, forming a spheroid as if there were no T cells at all.

## Methods

We performed *in silico* experiments in MATLAB with an agent-based model of T and cancer cells with stochastic behavior. These cells interact in three dimensions and are bounded by rules including cell division, migration, and killing, with underlying cytokine matrices determining their behavior. We created the model from scratch, using several matrices to track each cell type and its underlying behavior. The simulation layers are summarized in Figure 3a, which shows blue T cells and red cancer cells on the left and a 3-dimensional heat-map for the middle *x-y* slice of an example CCL2 matrix on the right. As seen in Figure 3b, cancer cells and T-cells are able to move within a radius around themselves in their respective matrices in three dimensions to any space unoccupied by other cells. Because each cancer cell and T cell acts independently every timestep, the model is agent-based. The full flow of the model as each timestep is processed is shown in Figure 3.

**Figure 3:**
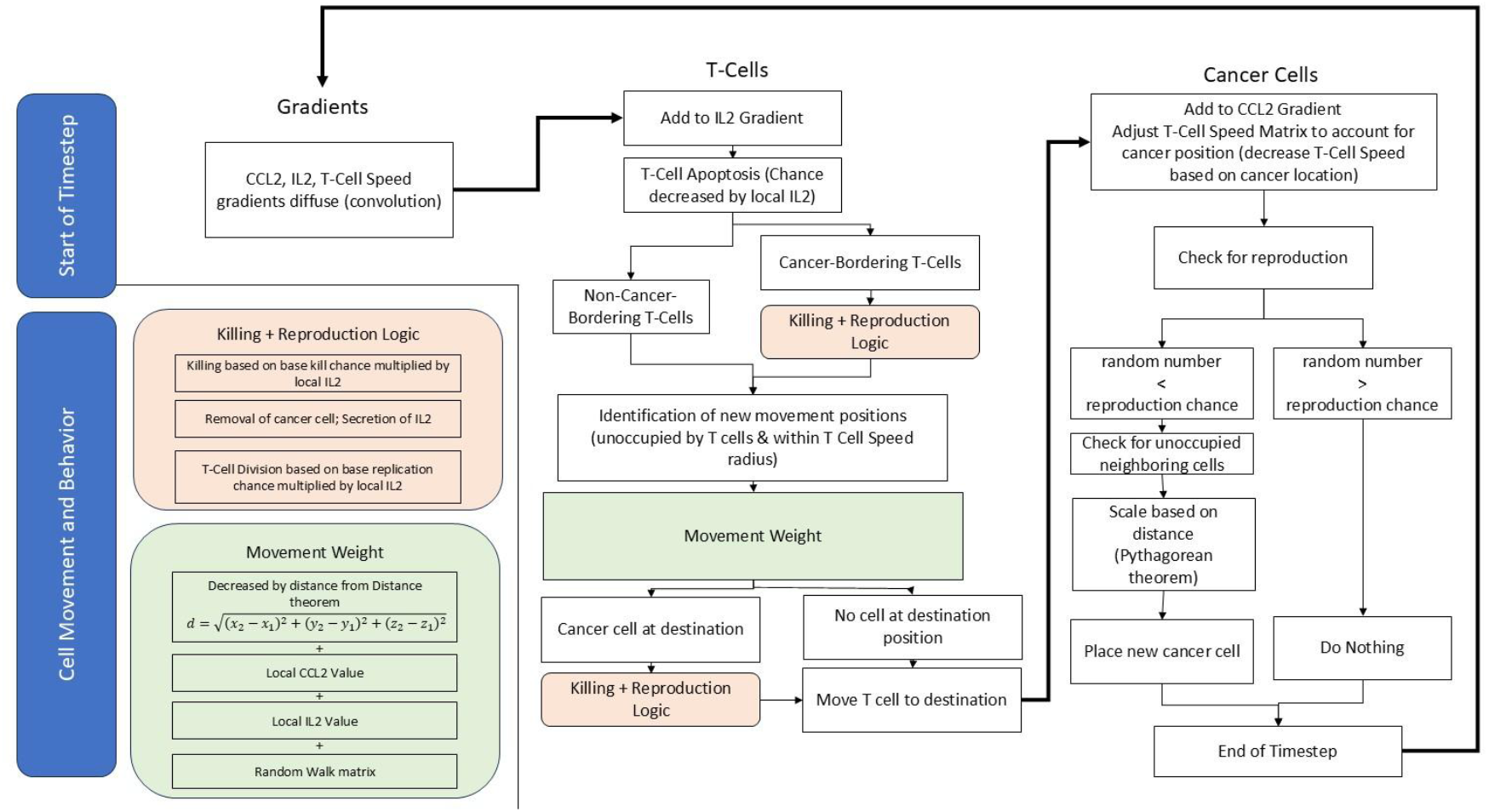
Simulation Flow

### Timescale and Size Scale

Each unit of simulation space is approximately 30 μm (the established cell size of breast cancer cells (14)) and each timestep is 10 minutes. Movement, replication, and killing speeds are scaled accordingly. Parameters and their corresponding established *in vitro* values are summarized in the table below:

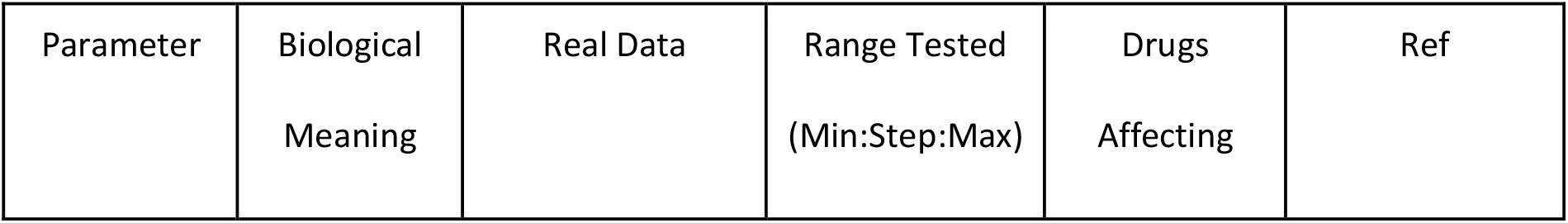
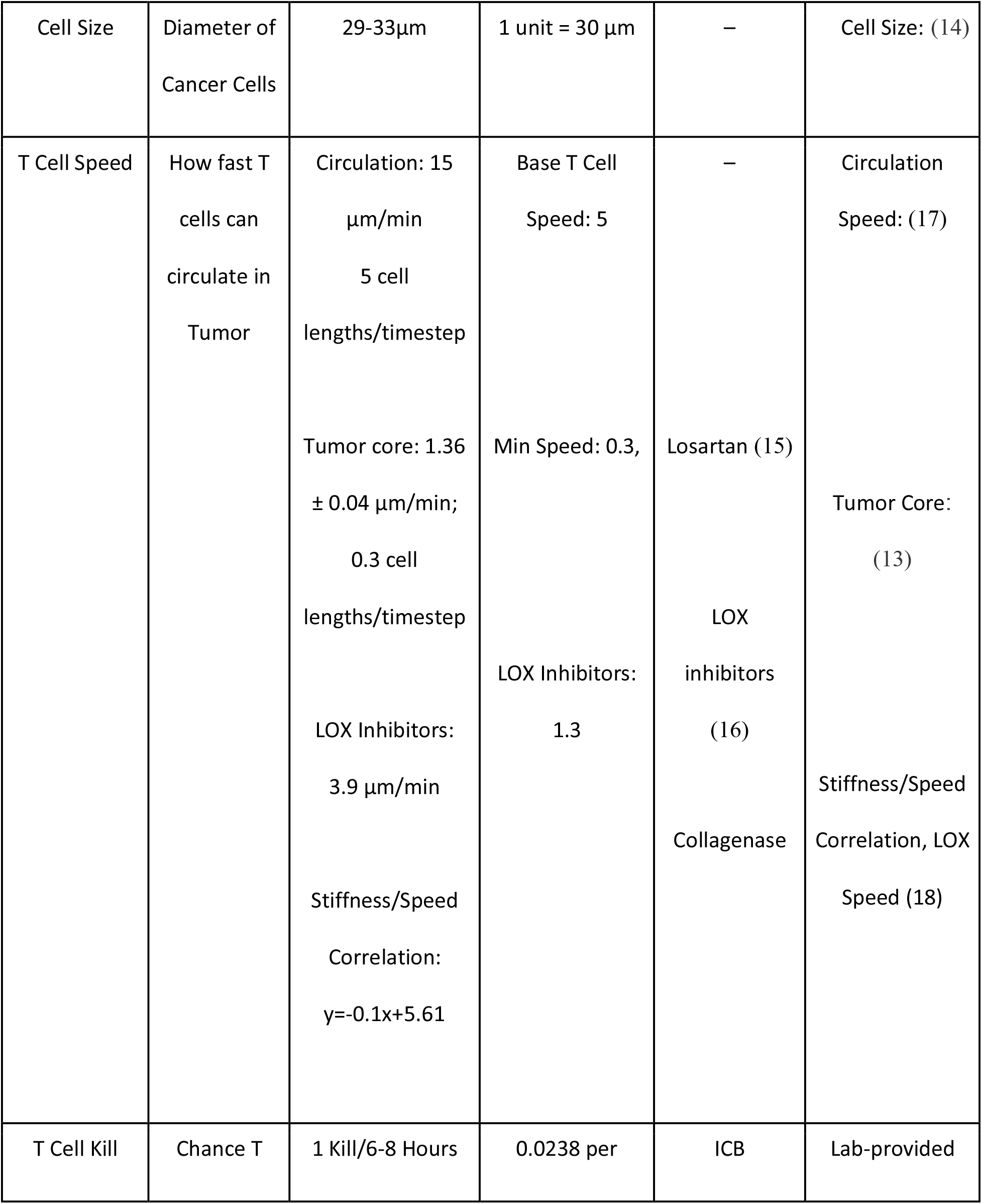
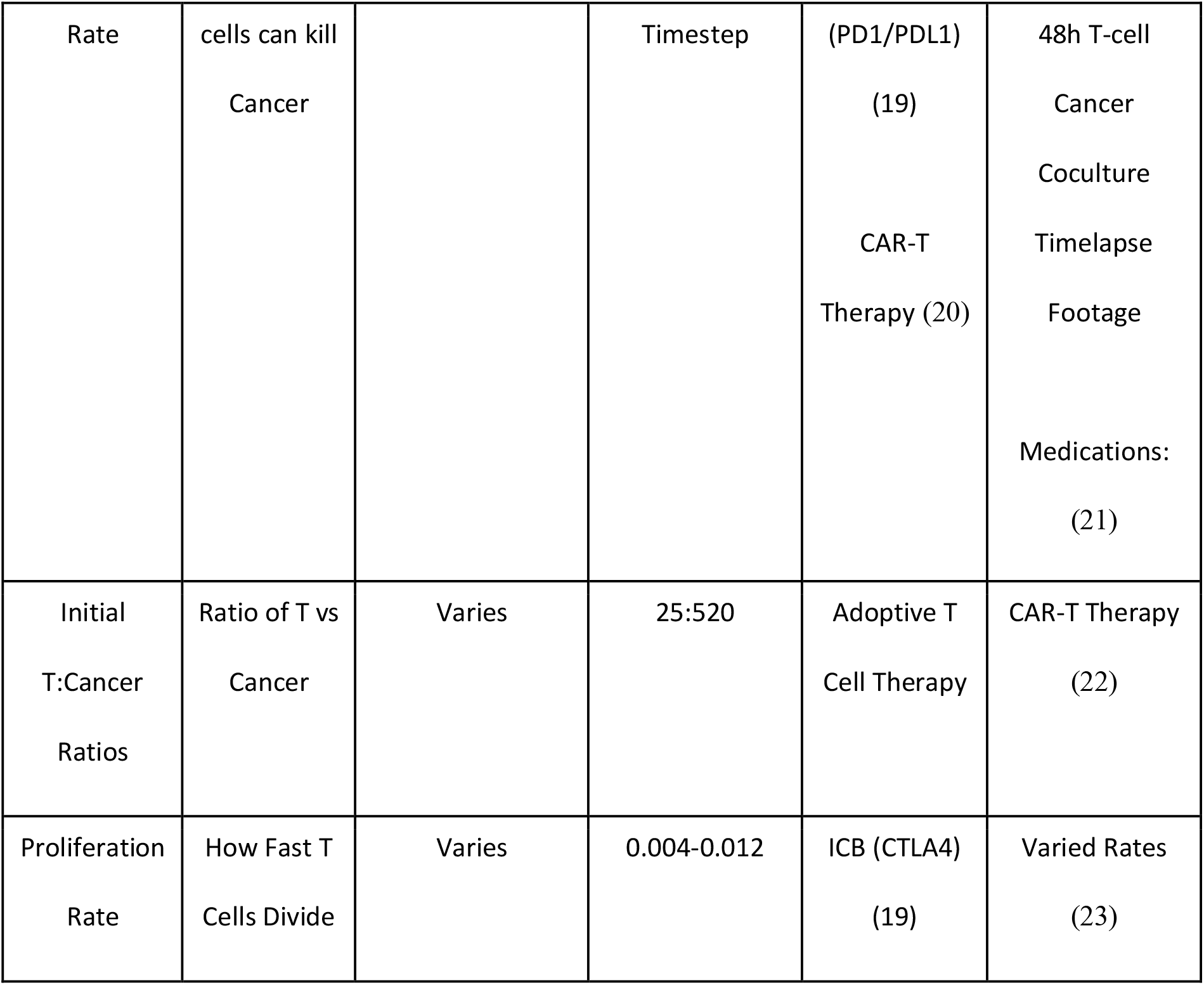

### Cell Matrices

We created two matrices, one for T-cells and one for cancer, that store the locations of each type of cell. Each matrix has the same size as the entire simulation space, with ones denoting the presence of a cell in that location and zeros denoting the absence of cells.

### Cytokine Matrices

We created several matrices to match the behavior of T-cells in tumors. Each matrix’s weight upon T-cell movement can be adjusted freely and independently. These matrices include:

#### IL-2 Matrix

We created a matrix storing the concentration of IL-2. IL-2 is produced at every time step by T-cells, and, using a convolution, each T-cell secretes a certain amount of IL-2 (`tCellIL2GradientAddition`) to the 3x3x3 grid of cells, including and immediately surrounding itself. Then, each time a cancer cell is killed, another amount (`tKillIL2Addition`) is released in the location of the now-dead cancer cell. IL-2 acts as a chemoattractant biasing T-cell movement and also increases T-cell activation and proliferation, especially in the tumor microenvironment, which in our simulation was represented in a higher chance of T-cells killing cancer based upon the local concentration of cancer and having a higher chance to reproduce when killing cancer. This matrix also diffuses (using another convolution) and degrades (using a linear multiplier) over time.

#### CCL2 Matrix

We then created a matrix storing the concentration of CCL2, another chemokine, across the simulation space. In reality, tumor cells both themselves produce and also induce other tumor-supporting cells in the tumor microenvironment to secrete CCL2, and the tumor cells in our simulation also produce CCL2, secreting it into the CCL2 matrix in the positions including and immediately surrounding themselves. The CCL2 acts as a chemoattractant to T-cells (as observed clinically), but does not have an effect on T-cell behavior other than chemotaxis. It also diffuses and degrades with each timestep.

#### Nutrient Matrix

Cancer cells are notoriously nutrient-deficient and have high metabolic rates, which also impede T-cell function. We created a nutrient matrix that is replenished whenever there are no cells, but is depleted by cancer cells and T-cells, with cancer cells depleting T-cells at a faster rate. This matrix diffuses over time, and acts as another influence upon T-cell behavior.

#### Random Walk Matrix

In addition to the two cytokine matrices, T cells are known to move semi-randomly, obeying behavior similar to Levy walks or Brownian motion depending on the strength of cell signaling surrounding them (24). This behavior is mimicked using a matrix full of random values between 0-1 which are scaled to be balanced against the aforementioned cytokine matrix. In this way, the T cells diffuse throughout the simulation space when there is a low concentration of the cytokines and undergo chemotaxis when the cytokine matrices have higher values.

### Cell Behavior

#### Cancer Cells

Cancer cells are generated in a ball at the beginning of the simulation. At each time step, cancer cells have a chance to reproduce into the cells adjacent to them, with a radius of 1.

Reproducing on the diagonal is proportionally less likely based on the total distance using the Pythagorean Theorem (dictated by the line `distances = sqrt(sum((validPositions - pos).^2, 2))`;). Cancer cells cannot grow into the same space (i.e. cancer cells in the center of the ball do not reproduce), and have no other behavior other than lowering the movement speed of T-cells (elaborated on later) and contributing to the CCL2 matrix. Their reproduction is tuned to the approximate doubling rate of breast cancer, which is once every 24 hours (144 timesteps) (14).

#### T-cells

##### T-cell Survival

T-cells are generated randomly in the space surrounding cancer. Each timestep, T-cells have a chance to survive. This chance is scaled by how much IL2 is in the area, with the survival chance approaching 100% as the local IL2 concentration approaches infinity. It is known that T-cells are less likely to transition to an anergic state or undergo apoptosis without a constant supply of costimulatory molecules, and undergo apoptosis without killing of target cells.

##### T-cell Mediated Cytotoxicity

T-cells have two chances per timestep to kill cancer: first, they can kill any cancer immediately surrounding them at the beginning of the timestep, and second, if they have selected to move to a space with a cancer cell, they have a chance to kill that cancer and take its position. Whenever a T-cell successfully eliminates a cancer cell, it releases IL2 and has a chance to reproduce into the now-empty cancer space, which is scaled linearly based on the IL2 concentration before the kill. Additionally, T-cells passively release a miniscule amount of IL2, which contributes to their clumping when cancer is not present (matching laboratory observation). The base chance of T-cell killing cancer was varied as a parameter, simulating therapeutic techniques which directly improve T-cell killing when interacting with cancer, such as PD-1/PDL1 Immune Checkpoint Blockade.

##### T-cell Proliferation

Whenever T-cells kill a cancer cell, they have a chance to reproduce into the space where the tumor was (if the killing took place before the T cell movement took place) or to reproduce into the space where the T cell was (thus yielding a T cell at its initial position and a new one at the killed tumor cell). This chance is linearly scaled by the local IL2 concentration. This parameter was also varied to simulate increased T-cell proliferation and activation, such as that achieved by CTLA4 Immune Checkpoint Blockade.

##### T-cell Movement

T-cells are able to move freely in a radius around themselves, except for when they move to a space occupied by other T-cells. The maximum radius is a preset, and at all locations, T-cells are initially able to move at that maximum radius (5 units/timestep, matching with *in vitro* observation of 15 μm/min for circulating T cells (17)). However, cancer cells take away an amount (called *ecmCancer*, since cancer is known to excrete excess protein which forms an extracellular matrix) which slows the speed T-cells can move at every time step when they approach the tumor spheroid. This speed matrix then “diffuses,” spreading out the impeding movement from the cancer. The speed matrix controls the range of new positions that T cells sample when considering new positions every timestep. For example, when the movement radius is 4.5, the T-cell has a 50% chance of being able to move between 5 units of itself, and a 50% chance of being able to move within 4 units of itself. The absolute hard minimum for the T-cell movement speed, and therefore the ease of T-cell movement within the tumor, was varied for the experiments. This behavior matches known treatments like Losartan (15) and LOX inhibitors (16), which reduce tumor stiffness and, in turn increase T cell mobility (18).

## Discussion and Future Directions

Clinically, αCTLA-4 therapy is known to maintain T-cell expansion (19), but in many cases, the therapy is unable to control the tumor. We recapitulated this behavior in our experiments (the 15% proliferation rate increase, although delaying the tumor growth, failed to regress the tumor). Our simulation identified that further increase of proliferation rate would be able to control the tumor, suggesting that methods pushing T cells towards a higher proliferation rate may exhibit improved tumor control. Interestingly, our model revealed that increasing the killing rate as the most effective method to enhance tumor control, a 17% increase is sufficient in controlling tumor growth at an early time point (Fig. 3, Fig. 4). These results from our model suggest that therapy that aims at increasing T cell killing rate, like CAR-T cells, may have stronger T cell killing enhancement even at minor changes. This recapitulates the clinical observation that CAR-T cells demonstrate significant improvement in treating hematological malignancy and some solid tumors, especially when coupled with treatments increasing their proliferation rate (25).

**Figure 4:**
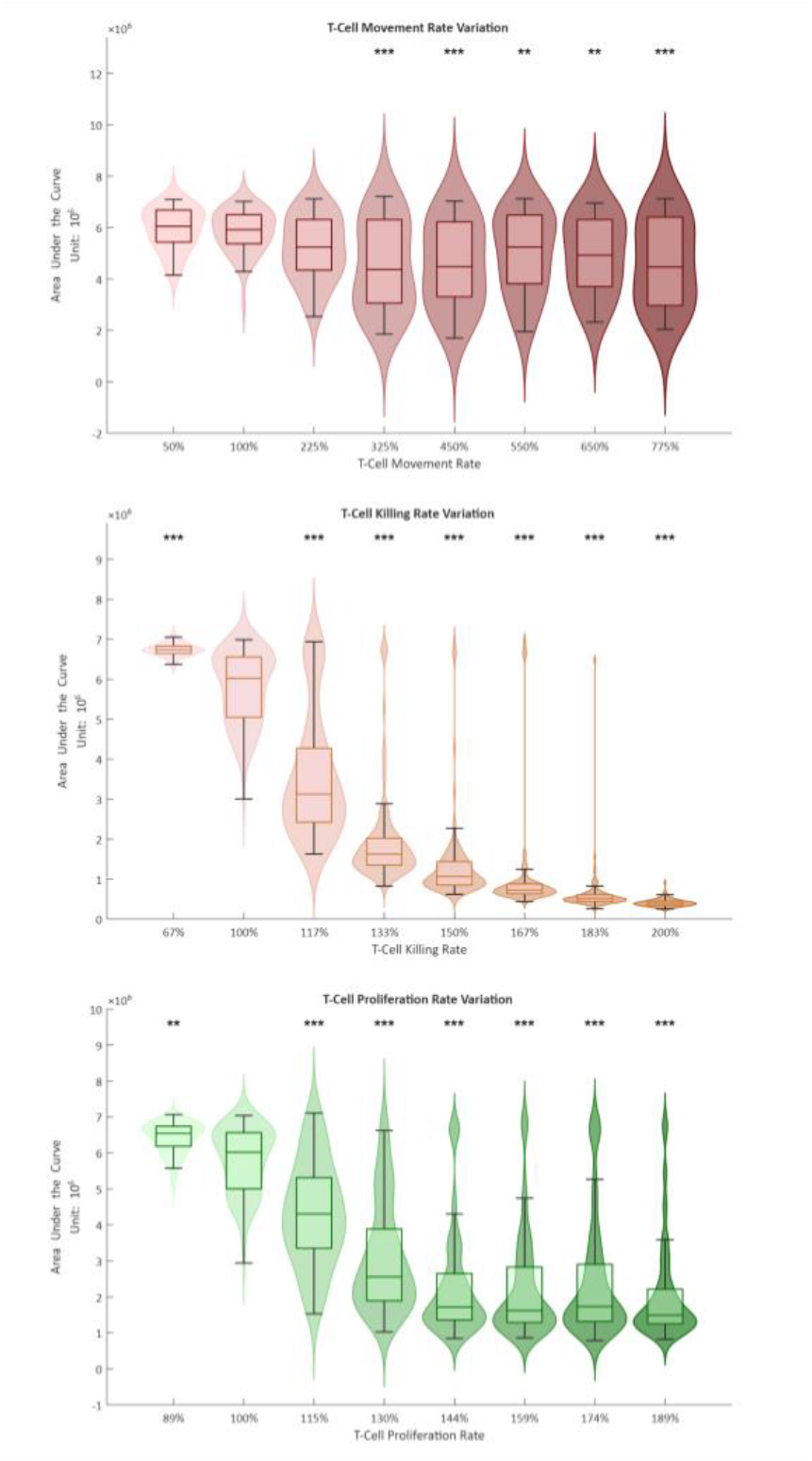
Area Under the Curve Plots by Condition

This simulation can be applied more broadly to optimize many other variables besides just T-cell killing, proliferation, and movement within the tumor. Additionally, the simulation can be tailored to match more cancers beyond triple negative breast cancer. The simulation can also be augmented to include mutations within the tumor and explore how the mutational burden of the cancer affects the immune response. More immune cell subtypes like M1 and M2 Macrophages as well as CD4+ T cells could be explored as well in simulation to augment the simulation’s scope.

## Acknowledgements

The authors would also like to extend an additional heartfelt thanks to Mr. Scott Balicki and Ms. Kathleen Bateman, who provided advice throughout the project and funding of the MATLAB software license. This project would not have been possible without their support and assistance.

